# Neural Networks model biological evolution of faithful epigenetic inheritance

**DOI:** 10.1101/2024.06.04.597202

**Authors:** B. N. Balakrishna Prabhu, Sibi Raj B. Pillai, Nithya Ramakrishnan

**Affiliations:** Institute of Bioinformatics and Applied Biotechnology, Bengaluru, India, 560100; Indian Institute of Technology Bombay, Mumbai, India - 400076

**Keywords:** Epigenetic Inheritance, Neural Networks, Histone Post-Translational Modifications

## Abstract

The layer of histone Post-Translational Modification (PTM) patterns, present above the DNA strand, forms an important epigenetic marker sequence which regulates gene expression. The specific pattern of histone PTMs in the region of chromatin housing the gene is critical for turning on/off the expression of the corresponding gene. During DNA replication in mitotic cells, the available evidence suggests that the histone PTMs from the mother chromatid are transferred uniformly at random among the two daughter chromatids. Parental epigenetic memory as well as interactions among multiple PTMs at the same histone facilitates the reconstruction of the PTM sequence at the daughter chromatids. We show that this biological marvel aided by the epigenetic memory has evolutionary analogs in the sense that it can be learnt by an appropriate extended neural network. We show through simulations that high fidelity reconstruction of the mother chromatin’s patterns for certain PTMs can be achieved by our network. This model can be enhanced to include several more interacting histone PTMs, elucidating the role of each. The proposed neural network can possibly be used in a multitude of biological applications related to gene expression regulation.

## I. Introduction

**H**ISTONE Post-Translational Modifications (PTMs) are one of the major epigenetic marks that regulate gene expression without altering the underlying genetic sequence [1]. Since the genetic sequence is identical in all cells of a given organism, it is the epigenetic marks which mainly dictate the functional properties of each cell. The PTM marks can be thought of as chemical functional groups attached to the protean spools (called histones) that help organize the DNA strand to form the chromatin structure, permitting selective access to the genes. A histone and its attached chemical marks together form a nucleosome, and they are distributed along the run of the DNA strand, found at an average distance of 200-250 DNA base-pairs apart. To maintain the smooth functioning of any group of cells, the PTMs are inherited across several cell generations. On the other hand, these marks are also impacted by external factors such as environment, diet, and stress [2]. The patterns of the PTMs across the promoter and gene-body play important roles in the activation or repression of genes [3]. While some histone PTMs (e.g. H3K27me3, where a trimethyl group is added to the 27th lysine on the tail of H3) silence the chromatin region in which they are present, a few others like H3K4me3, and H3K36e3 activate the underlying gene for transcription [4], [5].

When DNA replicates, each parental nucleosome is transferred to one of the daughter chromatids as a tetramer ((*H*3 − *H*4)_2_). It is postulated that the chromatid for each nucleosome placement is uniformly and independently chosen [6], [7]. A newly assembled nucleosome is placed on the other strand, thus conserving the total number of nucleosomes. Thus, about half of the parental histone tetramers are placed randomly in a daughter chromatid, this is akin to a discrete signal corrupted by an independent and identically distributed (i.i.d.) noise [8]. Thus, immediately after DNA replication, a daughter chromatid has two kinds of nucleosomes - one set that is directly inherited from the mother chromatin with its PTMs, and another set from a histone pool without any PTMs.

Certain histone PTMs such as H3K9me3 and H3K27me3 are essential for the inheritance for gene regulatory factors such as heterochromatin [9], [10]. It is therefore imperative that the distribution of histone PTMs in the parental chromatin is replicated with sufficient fidelity in the daughter chromatids, even if they have only half of the values.

To aid in the inheritance of the pattern of a histone PTM, histone chaperones and several other histone PTMs (that are antagonistic or synergistic in functionality) are known to interact with one another [10], [11]. The enzyme machinery and histone chaperones help in modifying the newly generated tetramers in the daughter chromatin based on the sequential pattern of modifications acquired from the parental chromatin [12]–[14].

Our previous work modelled the DNA replication process by a noisy communication channel, where the inheritance of a specific modification was aided by localized interactions amongst nucleosomes [15]. We then posed a question on whether the inheritance of a specific histone PTM such as H3K27me3 is aided by the presence of another histone PTM that is antagonistic to it (such as H3K36me3). If the answer to this is in the affirmative, then how much does the presence of an antagonistic mark improve the fidelity of the original mark? We explored the above questions using estimation theory principles in [16]. We investigate an alternate viewpoint in the current paper, can the solutions from the estimation theory framework evolve as a natural evolutionary solution. Towards this end, we train neural networks so as to learn and predict the pattern in the mother chromatin from the daughter chromatin. The network not only learns to estimate the mother pattern from the daughter sequence, but also can factor in the availability of additional information provided by an antagonistic sequence to improve the fidelity. We also show that the proposed neural network can be employed for predicting various statistical parameters related to the chromatin.

## II. Model

For a particular PTM, the binary digits 1 and 0 represent the presence and absence of that PTM at a nucleosome. Two PTMs are said to be antagonistic to each other when it is less likely to have have both of them simultaneously present on any nucleosome. Let us consider two antagonistic histone PTMs (say *A* and *B*), with PTM *A* being the principal modification of interest. Consider 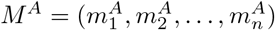 and 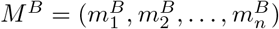 to be be the binary sequences representing the presence or absence of the histone PTMs *A* and *B* respectively across *n* nucleosomes in the mother chromatin. An example chromatin segment is shown in Fig. 1a. We model *M*^*A*^ as a first-order Markov chain similar to our previous work [15], whereas *M*^*B*^ is modelled as a Hidden Markov Model (HMM) driven by the Markov process *M*^*A*^. Using the Markov property, the conditional probability for *M*^*A*^ obeys

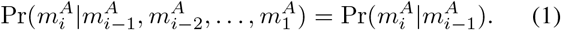

**Fig. 1.**
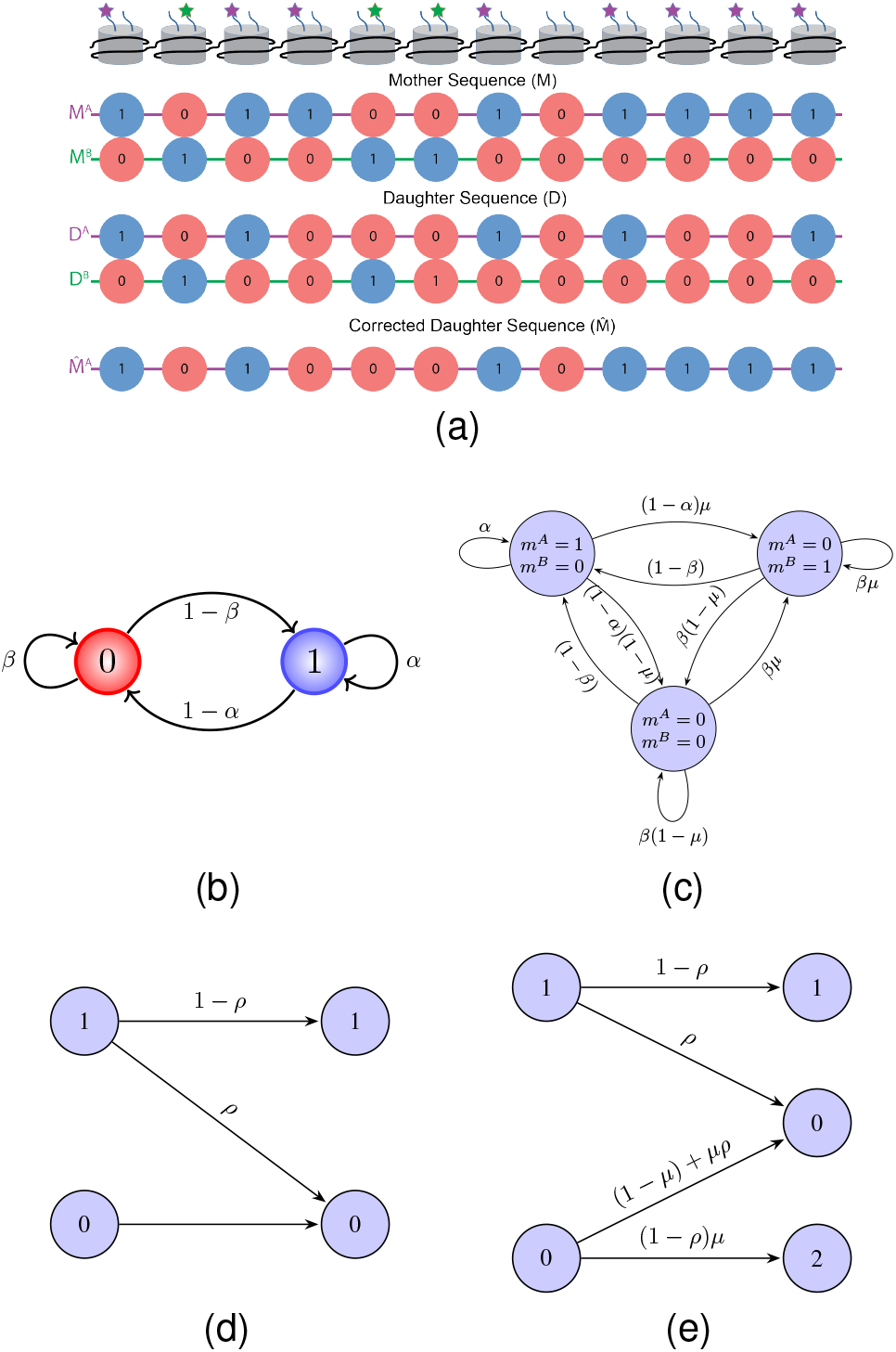
(a) The schematic representation of the antagonistic model where 1s and 0s represent the presence and absence of modifications. (b) State diagram for the case considering only PTM *A*. (c) State diagram for the case considering antagonistic modifications PTM *A* and PTM *B* assuming non-bivalent chromatin, i.e. 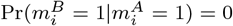. (d) DNA replication in the single modification case - modelled by a Z-Channel. (e) DNA replication in the antagonistic modification case - modelled by a Binary Erasure Channel.

**Fig. 2.**
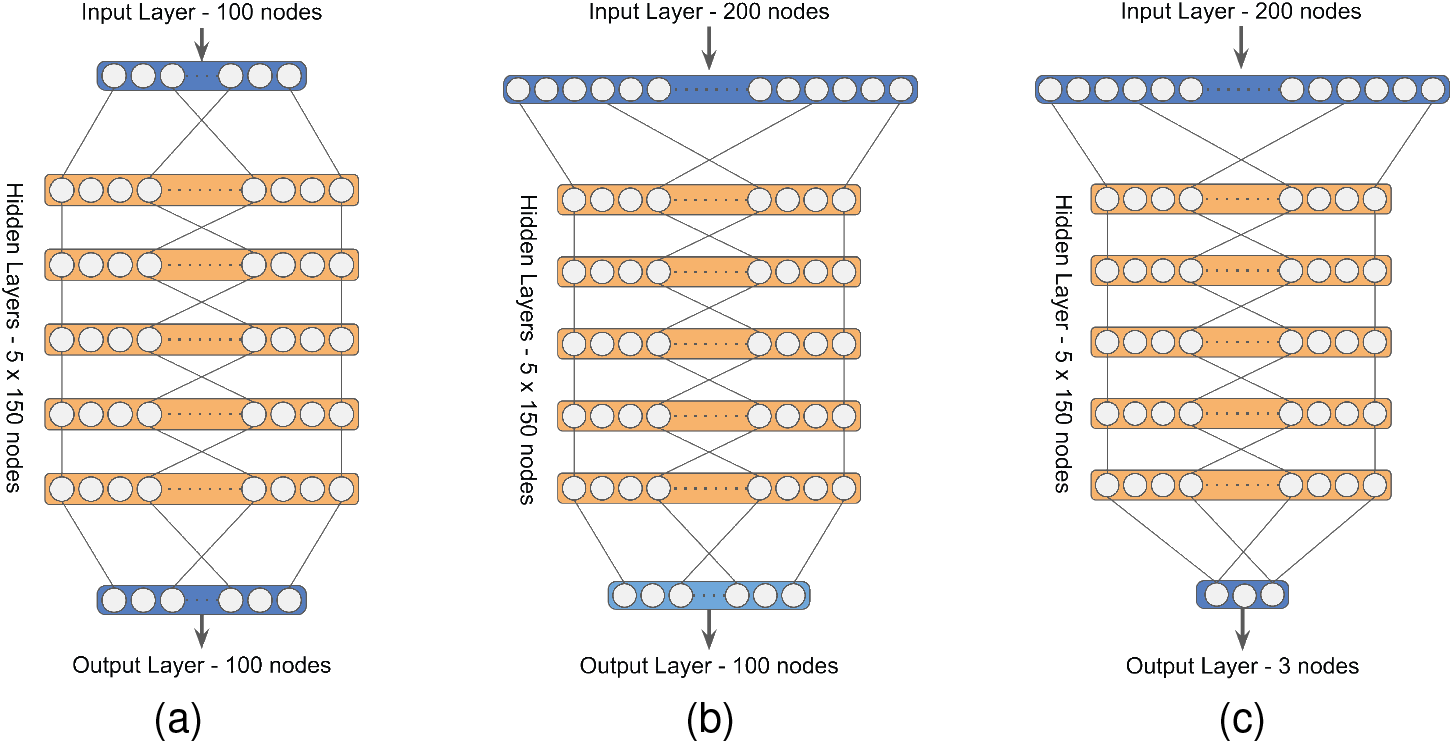
Neural Network Structures: (a) Network (targeted) for predicting *M*^*A*^ from *D*^*A*^, (b) Network (targeted and compound) for predicting *M*^*A*^ from *D*^*AB*^, (c) Network (compound) for prediction of *α, β*, and *µ* from *M*^*A*^

For the HMM *M*^*B*^, the value 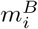 is conditionally independent of others, given 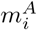.

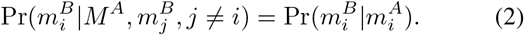

We represent the state transition diagram for histone PTM *A* alone in the mother chromatin in Fig. 1b. This is extended to include PTM *B* in Fig. 1c. The states 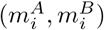 take values from the set {(0, 0), (0, 1), (1, 0)}. We make the assumption that the chromatin that we model for inheritance is not bivalent [17], i.e., we do not consider antagonistic modifications *A* and *B* to be present in the same nucleosome. Hence, (1, 1) is not included in the set of permissible states.

As can be seen from Fig. 1c, in addition to the transition probabilities *α* (probability of staying in a modified state) and *β* (probability of staying in an unmodified state), we define a new parameter *µ* as the probability of histone PTM *B* being present given that histone PTM *A* is absent in the same nucleosome. That is, 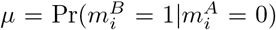 . Notice that 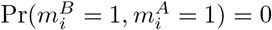 and 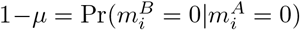.

As mentioned earlier, half of the histones are made available at the daughter chromatid uniformly at random. Let *D*^*AB*^ denote the sequence available at the daughter chromatid, where a de nuovo histone is assumed to have a state (0, 0). In Fig. 1e, we mark the output for the states (0, 0), (1, 0) and (0, 1) as 0, 1 and 2 respectively. The objective of the neural network is to estimate the mother sequence from the observations at a daughter chromatid. To compute the fidelity of reconstruction, we define the Bit Error Rate (BER) corresponding to *M*^*A*^ as the fraction of nucleosomes where the original and estimated mother sequence 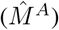 differ with respect to PTM *A*, i.e.,

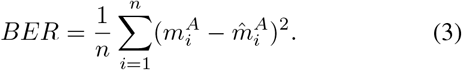

### A. Neural Network for Histone PTMs

For a single PTM sequence following the Markov model [15], it was shown recently that the inheritance of a histone PTM from the mother to the daughter chromatin has high fidelity when it follows a threshold-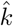 filling algorithm in certain regions of the chromatin (long islands of 1s interspersed by long islands of 0s). This was then extended to accommodate the presence of antagonistic modifications [16]. These works employed the Sequential Maximum A Posteriori Probablity (SMAP) maximizing Viterbi algorithm to decode the sequences and arrive at the respective inferences.

In the area of epigenetic inheritance, the first question was whether evolutionarily one can mimic the optimal algorithms such as threshold-k filling - which we proved to be optimal for SMAP decoding by analytical techniques, when antagonistic PTMs are absent. While the threshold filling has to incorporate the specific transition probabilities to identify the threshold, a well trained neural network may provide a more universal decoding strategy, across all values of *α* and *β*. Additionally in the presence of interacting modifications (such as antagonistic modifications), how is the decoding impacted? Quite interestingly, an appropriately trained neural network can not only mimic the patterns of inheritance in specific regimes but also provide a more universal decoding strategy.

### B. Neural Network Designs

Based on the range of the HMM parameters passed to the network, we consider the following types: (i) targeted network, specific to a (*α, β, µ*) tuple and (ii) compound networks, trained over all ranges of (*α, β, µ*) values.

We use these network architectures to perform the following experiments:

- Predict *M*^*A*^ from *D*^*A*^ - (targeted network)
- Predict *M*^*A*^ from *D*^*AB*^ - (targeted and compound network)
- Predict *α, β*, and *µ* from *M*^*AB*^ - (compound network) The parameters for the neural network architectures are listed in Table. I.

**TABLE 1.** Parameters for network experiments.

| Neural Network Parameters |  |  |  |  |
| --- | --- | --- | --- | --- |
| Parameter | $M^A$ from $D^A$ | $M^A$ from $D^{AB}$ | $M^A$ from $D^{AB}$ | $\alpha, \beta, \mu$ from $M^{AB}$ |
| Network Type | Targeted | Targeted | Compound | Compound |
| Random Seed | 42 | 42 | 42 | 42 |
| Sequence Length | 100 | 100 | 100 | 100 |
| Training Samples ($S_{\text{simple}}$) | 100,000 | 100,000 | 100,000 | 10,000 |
| Testing Samples | 30,000 | 30,000 | 30,000 | 3,000 |
| Loss Function Type | MSE | MSE | MSE | MSE |
| Non-Linear Activation | ReLU | ReLU | ReLU | ReLU |
| Input Size | 100 | 200 | 200 | 200 |
| Number of Layers | 5 | 5 | 5 | 5 |
| Hidden Size | 150 | 150 | 150 | 150 |
| Output Size | 100 | 100 | 100 | 3 |
| Learning Rate | 0.001 | 0.001 | 0.001 | 0.001 |
| Epochs ($E_{\text{simple}}$) | 10,000 | 10,000 | 20,000 | 10,000 |
| Batch Size | 100,000 | 100,000 | 100,000 | 10,000 |

## III. Results

We first analyze the results with the targeted networks. Simulated daughter sequences *D*^*A*^ each of length 100 are passed as input to be trained to produce the reconstructed mother 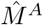. The output sequences are compared with the actual mother sequence (*M*^*A*^) to compute the average *BER* across 30,000 such sequences. One neural network was trained for each (*α, β*) pair.

We then introduce the antagonistic daughter sequence *D*^*B*^ along with *D*^*A*^ to the targeted networks to check if there is an improvement in the fidelity of reconstruction of *M*^*A*^. The input to each network is doubled from the single modification case. As can be seen from Fig. 3b the presence of the antagonistic daughter sequence (for *µ* = 0.5) improves the fidelity (reduces the *BER*) of the estimated 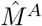. This can also be observed through the Hinton plot in Fig. 3c.

Next, we look at the results for the compound network. Here, we train the network with 100,000 sequences with parameters (*α, β*) ∈ [0.0, 1.0) and *µ* ∈ [0.0, 1.0) using 20000 epochs. We obtained the average *BER* across 1000 samples for each (*α, β, µ*) tuple. In Fig. 4a and 4b, we plot the heatmaps obtained using the compound network for the cases of *µ* = 0 (no-antagonism case) and *µ* = 0.5 respectively. From these figures and Fig. 4c, one can observe that, with the presence of antagonism, the *BER* decreases, similar to the targeted network case. Refer to Fig. A1 in Appendix 1 for the heatmaps and Hinton plots corresponding to other values of *µ*.

In Fig. 4d, we plot the variation of the average *BER* for a few selected (*α, β*) pairs with *µ*. It is evident from this plot, that antagonism helps to improve the fidelity of inheritance of the primary modification.

The Neural Network depicted in Fig. 2c learns and predicts the parameters of the HMM model (*α, β* and *µ*) from simulated mother sequences (*M*^*AB*^). We plot the Pearson’s correlation between the actual parameter values of the sequences and the predicted parameters in Fig 5 (a,b,c). As can be seen from the figures, the predicted values correlate with the parameter values of the input sequences. The results of the HMM predictor network validates our model for epigenetic inheritance.

**Fig. 3.**
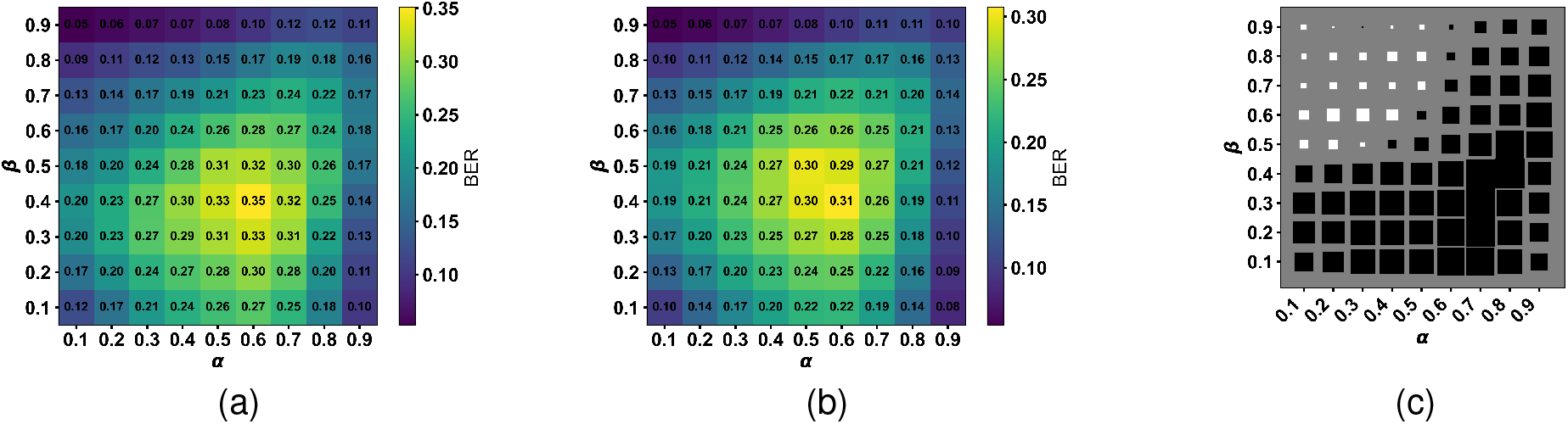
Simulation results for different *α, β* combinations: The figures (A, B) show the average *BER* values for the different (*α, β*) combinations for the non-antagonistic (*µ* = 0.0) and antagonistic (*µ* = 0.5) cases with the targeted networks. (C) Shows the corresponding Hinton plots highlighting the difference between *BER* of the two. In the Hinton plot, a black square shows that antagonism is providing a lower *BER* than the no-antagonism counterpart for that (*α, β*). In the Hinton plots, the largest square has a magnitude of |0.0596|.

**Fig. 4.**
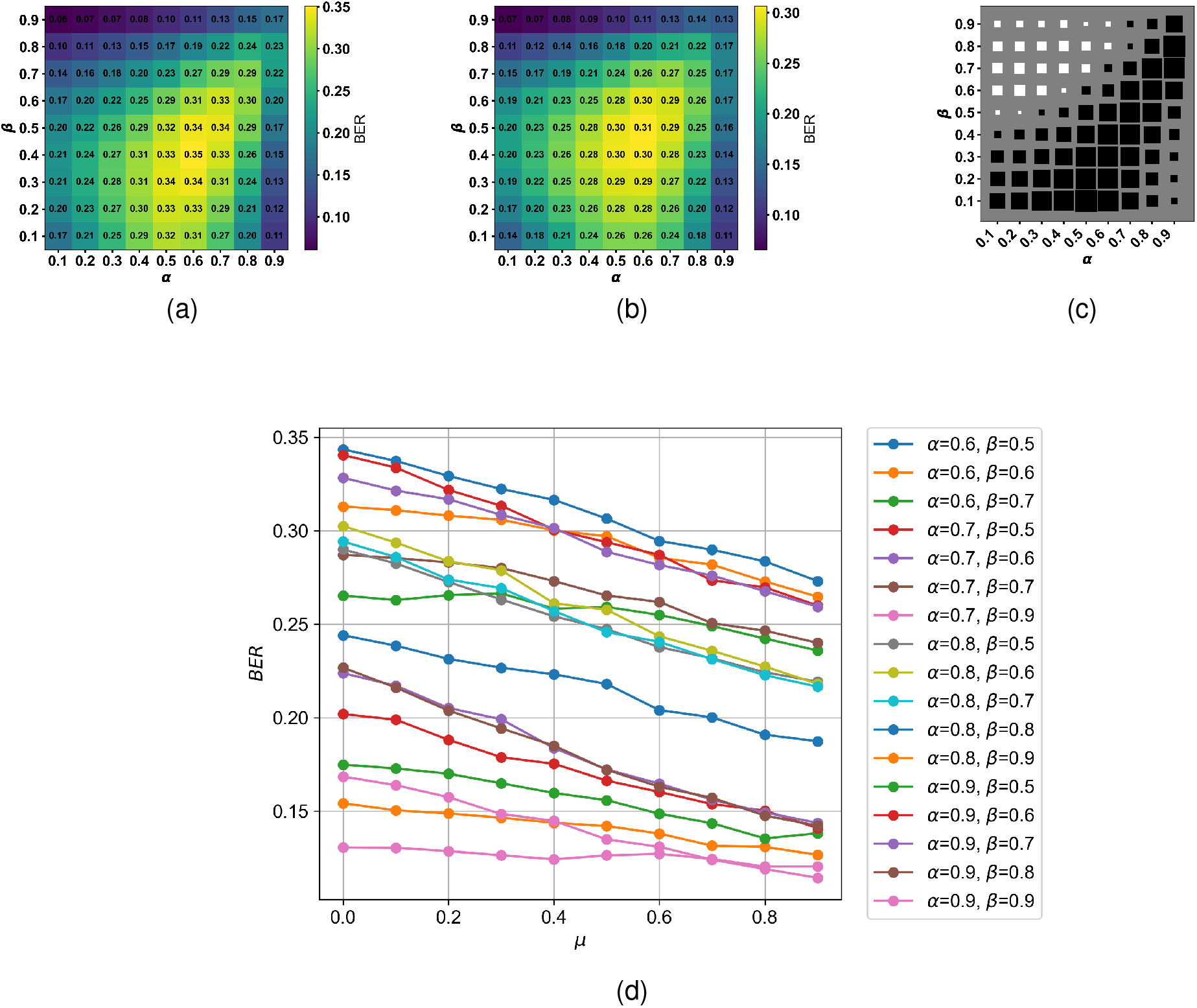
Heatmaps from simulations using a compound neural network trained over *α, β, µ* ∈ [0, 1] for (a) *µ* = 0.0. (b) *µ* = 0.5. (c) Hinton plots for comparison of *BER* between *µ* = 0.0 and *µ* = 0.5 for the compound neural network with the largest square having a magnitude of |0.0576|. (d) Simulation results from the compound neural network trained over *α, β, µ* ∈ [0, 1] show how an increase in antagonism reduces the *BER* for different pairs of (*α, β*).

**Fig. 5.**
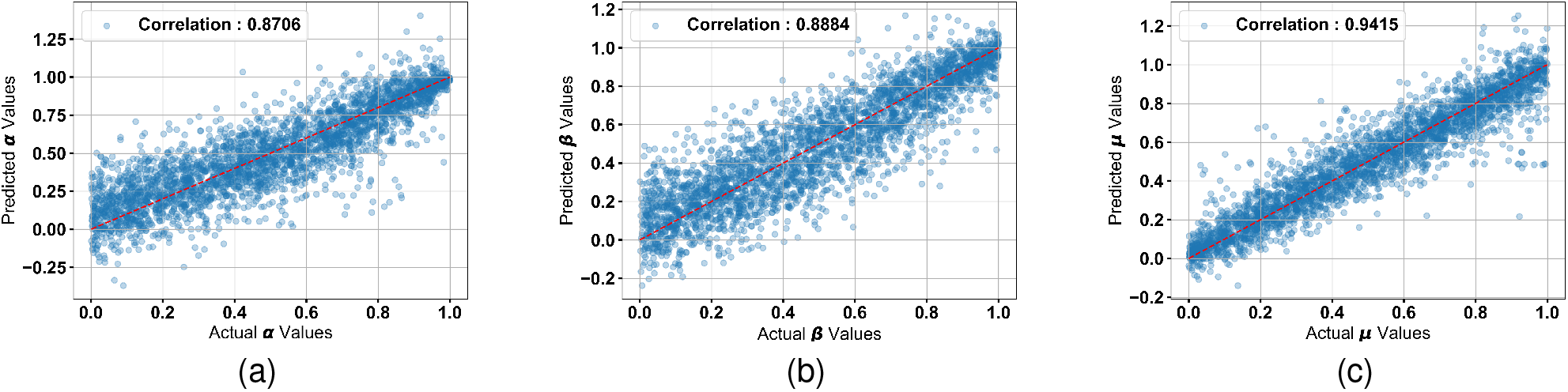
Plots showing the Pearson’s correlation between the predicted and actual values for (a) *α*, (b) *β*, and (c) *µ* using the HMM parameter predicting neural network.

## IV. Discussion and future work

Inheritance of epigenetic information is a complex biological process that depends on a plethora of factors such as the enzyme machinery that adds/removes modifications, histone chaperones and other proteins [7]. Particularly, the inheritance of the patterns of the histone PTMs is significant since it has to match the mother chromatin’s pattern early in the cell cycle [18]. In many cases, the inheritance of related epigenetic phenomena such as DNA methylation play a critical role in aiding the inheritance of histone PTMs [19]. Additionally, the histone PTMs are known to interact with one other and contribute to the gene regulation jointly as a histone code [20]. This complex set of factors can be efficiently modelled by a neural network - we show the fundamental case of a single modification and the case of antagonistic modifications in this paper.

Several statistical and probabilistic models have been proposed to explain the inheritance of epigenetic marks. Skjegstad *et al* proposed a 3D polymer configuration model in which they postulate that nucleosomes bind to one another in 3D configurations that affect phenomena such as epigenetic memory and bistability [21]. They employ partial confinement to explain the burst-like nature of certain epigenetic switches. A stochastic model was proposed by Dodd *et al* [22], in which they postulate that cooperation between neighbouring nucleosomes leads to the spreading of modifications over distant regions. Using experimental data from fission yeast, the authors in [23] suggest that heterochromatin propagates in bursts and not linearly. Alabert *et al* [24] experiment with two different models (global and domain) and suggest that the domain model (in which methylation states are found in distinct domains) is more likely to explain the antagonism between H3K36me3 and H3K27me3 in mESCs. These past works have not employed Neural Networks to model the spreading and inheritance of histone modifications.

Neural networks have been used to predict gene expression based on histone modification data in the past. For instance, DeepChrome [25] is a deep CNN that predicts gene expression based on five histone modifications from the REMC database [26]. The authors later used a Deep Learning model with attention mechanisms to improve the prediction in AttentiveChrome [27]. More recently, in [28], the authors use Transfer Learning to predict gene expression from the histone PTM database for cross-cell lines. There have also been a plethora of tools such as EP-DNN which is a Deep Neural Network for predicting enhancer signatures from chromatin features [29]. However, there have not been any neural network models for modeling and predicting epigenetic inheritance so far.

We compare the Neural Network results for epigenetic inheritance with our previous analytical approach (SMAP-based decoding). We find that the results are comparable, with the analytical case doing slightly better for certain ranges of HMM parameters (Fig A2 in Appendix). However, if we were to extend this study to more histone modifications (those that are antagonistic or synergistic), the analytical case may become increasingly complex to generalize, whereas the proposed extended neural network may still be able to learn the patterns and interactions between the modifications easily.

It should also be noted that we assume that the daughter sequence has been obtained with the parameter *ρ* set to 0.5 (symmetric inheritance). In biological systems, there may be certain types of cells where this may not be true [30]. It would be interesting to see how the mother sequence’s pattern is reproduced from the case of disadvantaged daughter sequences using the neural networks approach. These are some of the works for the future.

## Acknowledgments

This study is supported by the Dept. of Electronics, IT, BT and S&T, Government of Karnataka, India.

---

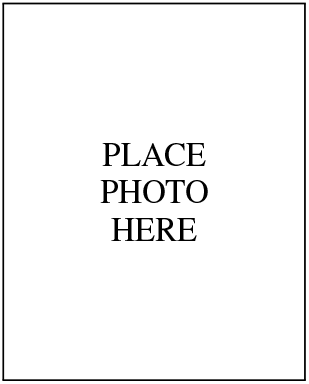

**B. N. Balakrishna Prabhu** received his Integrated Master of Science degree in Physics from Cochin University of Science and Technology in 2023. He is currently a Research Assistant at the Institute of Bioinformatics and Applied Biotechnology in Bangalore. His research interests include physics, computer science, evolutionary game theory, complex systems, and non-linear dynamics. Driven by a passion for lifelong learning, he is actively pursuing an academic career and is working towards a PhD.

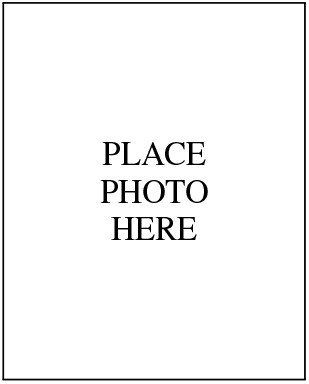

**Sibi Raj B. Pillai** received the PhD in computer science and communication systems from Ecole Poly-technique Fédérale de Lausanne (EPFL), Switzerland, in July 2007. He was a Research Fellow with the University of Melbourne from October 2007 to April 2009, and ever since a faculty member at the Department of Electrical Engineering, Indian Institute of Technology Bombay. His research interests are in network information theory, feedback communications, cross-layer scheduling, biological information inheritance, and radar signal processing.

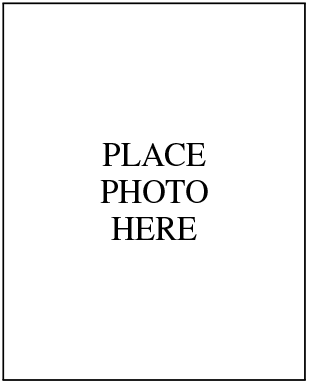

**Nithya Ramakrishnan** received her PhD from Indian Institute of Technology Delhi in the area of Computational Biology and Bioinformatics, from the Department of Electrical Engineering. She did a post-doctoral term at the Indian Institute of Technology Bombay in the Department of Biosciences and Bioengineering, followed by another post-doctoral term at Dalhousie University, Canada. Her interests include Machine Learning for multiomic data analysis and probabilistic modelling of biological inheritance. Prior to her research career, she worked in the software industry in the areas of cloud computing for over a decade. She joined the IBAB Faculty in early 2023.

## References

[1] C. L. Peterson and M.-A. Laniel, “Histones and histone modifications,” Current Biology, vol. 14, no. 14, pp. R546–R551, 2004.

[2] A. D. Goldberg, C. D. Allis, and E. Bernstein, “Epigenetics: a landscape takes shape,” Cell, vol. 128, no. 4, pp. 635–638, 2007.

[3] C. D. Allis, S. L. Berger, J. Cote, S. Dent, T. Jenuwien, T. Kouzarides, L. Pillus, D. Reinberg, Y. Shi, R. Shiekhattar et al., “New nomenclature for chromatin-modifying enzymes,” Cell, vol. 131, no. 4, pp. 633–636, 2007.

[4] H. Santos-Rosa, R. Schneider, A. J. Bannister, J. Sherriff, B. E. Bernstein, N. T. Emre, S. L. Schreiber, J. Mellor, and T. Kouzarides, “Active genes are tri-methylated at k4 of histone h3,” Nature, vol. 419, no. 6905, pp. 407–411, 2002.

[5] D. K. Pokholok, C. T. Harbison, S. Levine, M. Cole, N. M. Hannett, T. I. Lee, G. W. Bell, K. Walker, P. A. Rolfe, E. Herbolsheimer et al., “Genome-wide map of nucleosome acetylation and methylation in yeast,” Cell, vol. 122, no. 4, pp. 517–527, 2005.

[6] I. M. Hall, G. D. Shankaranarayana, K.-i. Noma, N. Ayoub, A. Cohen, and S. I. Grewal, “Establishment and maintenance of a heterochromatin domain,” Science, vol. 297, no. 5590, pp. 2232–2237, 2002.

[7] A. V. Probst, E. Dunleavy, and G. Almouzni, “Epigenetic inheritance during the cell cycle,” Nature reviews Molecular cell biology, vol. 10, no. 3, pp. 192–206, 2009.

[8] C. Alabert, T. K. Barth, N. Reverón-Gómez, S. Sidoli, A. Schmidt, O. N. Jensen, A. Imhof, and A. Groth, “Two distinct modes for propagation of histone ptms across the cell cycle,” Genes & development, vol. 29, no. 6, pp. 585–590, 2015.

[9] A. R. Cutter DiPiazza, N. Taneja, J. Dhakshnamoorthy, D. Wheeler, S. Holla, and S. I. Grewal, “Spreading and epigenetic inheritance of heterochromatin require a critical density of histone h3 lysine 9 tri-methylation,” Proceedings of the National Academy of Sciences, vol. 118, no. 22, p. e2100699118, 2021.

[10] T. M. Escobar, A. Loyola, and D. Reinberg, “Parental nucleosome segregation and the inheritance of cellular identity,” Nature Reviews Genetics, vol. 22, no. 6, pp. 379–392, 2021.

[11] A. Stirpe, N. Guidotti, S. J. Northall, S. Kilic, A. Hainard, O. Vadas, B. Fierz, and T. Schalch, “Suv39 set domains mediate crosstalk of heterochromatic histone marks,” Elife, vol. 10, p. e62682, 2021.

[12] G. Schlissel and J. Rine, “The nucleosome core particle remembers its position through dna replication and rna transcription,” Proceedings of the National Academy of Sciences, vol. 116, no. 41, pp. 20 605–20 611, 2019.

[13] K. R. Stewart-Morgan, N. Petryk, and A. Groth, “Chromatin replication and epigenetic cell memory,” Nature Cell Biology, vol. 22, no. 4, pp. 361–371, 2020.

[14] A. Corpet and G. Almouzni, “Making copies of chromatin: the challenge of nucleosomal organization and epigenetic information,” Trends in cell biology, vol. 19, no. 1, pp. 29–41, 2009.

[15] N. Ramakrishnan, S. R. B. Pillai, and R. Padinhateeri, “High fidelity epigenetic inheritance: Information theoretic model predicts threshold filling of histone modifications post replication,” PLOS Computational Biology, vol. 18, no. 2, p. e1009861, 2022.

[16] B. B. Prabhu, S. R. B. Pillai, and N. Ramakrishnan, “Antagonistic histone post-translational modifications improve the fidelity of epigenetic inheritance-a bayesian perspective,” bioRxiv, pp. 2024–05, 2024.

[17] K. Sneppen and L. Ringrose, “Theoretical analysis of polycomb-trithorax systems predicts that poised chromatin is bistable and not bivalent,” Nature communications, vol. 10, no. 1, p. 2133, 2019.

[18] T. M. Escobar, O. Oksuz, R. Saldaña-Meyer, N. Descostes, R. Bonasio, and D. Reinberg, “Active and repressed chromatin domains exhibit distinct nucleosome segregation during dna replication,” Cell, vol. 179, no. 4, pp. 953–963, 2019.

[19] A. Groth, W. Rocha, A. Verreault, and G. Almouzni, “Chromatin challenges during dna replication and repair,” Cell, vol. 128, no. 4, pp. 721–733, 2007.

[20] T. Jenuwein and C. D. Allis, “Translating the histone code,” Science, vol. 293, no. 5532, pp. 1074–1080, 2001.

[21] L. E. J. Skjegstad, J. F. Nickels, K. Sneppen, and J. B. Kirkegaard, “Epigenetic switching with asymmetric bridging interactions,” Biophysical Journal, vol. 122, no. 12, pp. 2421–2429, 2023.

[22] I. B. Dodd, M. A. Micheelsen, K. Sneppen, and G. Thon, “Theoretical analysis of epigenetic cell memory by nucleosome modification,” Cell, vol. 129, no. 4, pp. 813–822, 2007.

[23] J. F. Nickels, A. K. Edwards, S. J. Charlton, A. M. Mortensen, S. C. L. Hougaard, A. Trusina, K. Sneppen, and G. Thon, “Establishment of heterochromatin in domain-size-dependent bursts,” Proceedings of the National Academy of Sciences, vol. 118, no. 15, p. e2022887118, 2021.

[24] C. Alabert, C. Loos, M. Voelker-Albert, S. Graziano, I. Forné, N. Reveron-Gomez, L. Schuh, J. Hasenauer, C. Marr, A. Imhof et al., “Domain model explains propagation dynamics and stability of histone h3k27 and h3k36 methylation landscapes,” Cell reports, vol. 30, no. 4, pp. 1223–1234, 2020.

[25] R. Singh, J. Lanchantin, G. Robins, and Y. Qi, “Deepchrome: deep-learning for predicting gene expression from histone modifications,” Bioinformatics, vol. 32, no. 17, pp. i639–i648, 2016.

[26] A. Kundaje, W. Meuleman, J. Ernst, M. Bilenky, A. Yen, A. Heravi-Moussavi, P. Kheradpour, Z. Zhang, J. Wang, M. J. Ziller et al., “Integrative analysis of 111 reference human epigenomes,” Nature, vol. 518, no. 7539, pp. 317–330, 2015.

[27] R. Singh, J. Lanchantin, A. Sekhon, and Y. Qi, “Attend and predict: Understanding gene regulation by selective attention on chromatin,” Advances in neural information processing systems, vol. 30, 2017.

[28] Y. Chen, M. Xie, and J. Wen, “Predicting gene expression from histone modifications with self-attention based neural networks and transfer learning,” Frontiers in Genetics, vol. 13, p. 1081842, 2022.

[29] S. G. Kim, M. Harwani, A. Grama, and S. Chaterji, “Ep-dnn: a deep neural network-based global enhancer prediction algorithm,” Scientific reports, vol. 6, no. 1, p. 38433, 2016.

[30] J. Xie, M. Wooten, V. Tran, and X. Chen, “Breaking symmetry-asymmetric histone inheritance in stem cells,” Trends in cell biology, vol. 27, no. 7, pp. 527–540, 2017.

