## Appendix for "Neural Networks model biological evolution of faithful epigenetic inheritance"

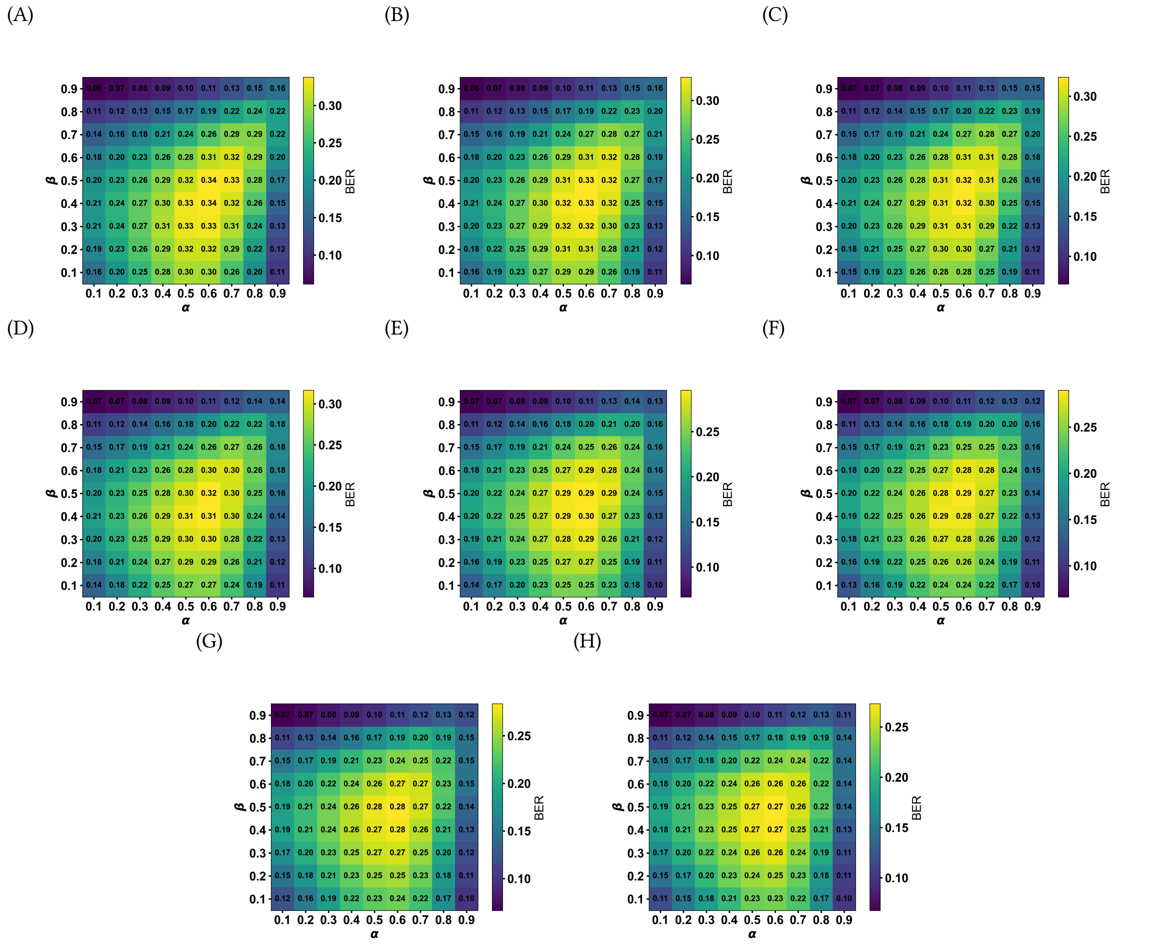


Fig. A1. Annotated heatmaps from simulations using a single neural network trained over $\alpha, \beta, \mu \in [0, 1]$. (a) $\mu = 0.1$. (b) $\mu = 0.2$. (c) $\mu = 0.3$. (d) $\mu = 0.4$. (e) $\mu = 0.6$. (f) $\mu = 0.7$. (g) $\mu = 0.8$. (h) $\mu = 0.9$.

Authors' addresses: B. N. Balakrishna Prabhu, Institute of Bioinformatics and Applied Biotechnology, Electronics City Phase I, Bengaluru, India, 560100,; Sibi Raj B. Pillai, Indian Institute of Technology Bombay, Powai, Mumbai, India, 400088,; Nithya Ramakrishnan, Institute of Bioinformatics and Applied Biotechnology, Electronics City Phase I, Bengaluru, India, 560100,.

(A) (B)

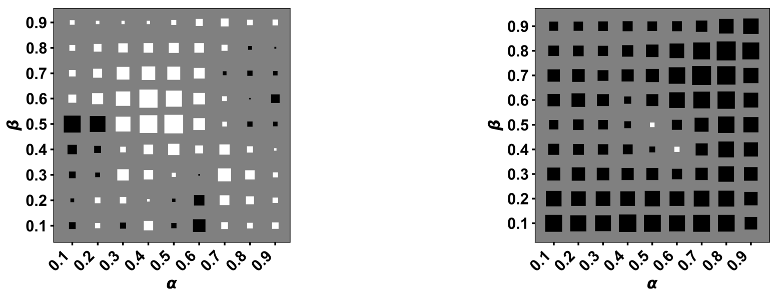


Fig. A2. Hinton plots highlighting the difference between *BER* between Viterbi Decoding vs Neural Network decoding. In the Hinton plot, a black square shows that Neural Network is providing a lower *BER* than the Viterbi decoding counterpart for that $(\alpha, \beta, \mu)$. In the Hinton plots, the largest square has a magnitude of $|0.1|$. (A) Targeted Network vs Viterbi for $\mu = 0.5$ black showing targeted network performing better, (B) Targeted Network vs compound network for $\mu = 0.5$ black showing targeted network performing better
